# Trait motivation is associated with Fusiform face area Morphometry: Evidence from a Chinese Youth Sample

**DOI:** 10.1101/2025.10.03.678225

**Authors:** Hangshek Lau, Yiying Song, Shan Xu, Jia Liu

## Abstract

Trait motivation is fundamental in shaping human behaviors. Previous studies have primarily focused on their impact on affective and motivational processing, with their role in perceptual processes less investigated. The present study takes face perception, a crucial and well-studied perceptual process, as a representative specimen to examine the perceptual effect of trait motivation. We investigated whether the behavioral activation system (BAS) and the behavioral inhibition system (BIS) were associated with structural characteristics of the inferior temporal face-selective regions as well as face recognition performance. With a sample of Chinese young adults (N = 264), voxel-based morphometry revealed that BIS scores correlated with greater gray matter volume in the fusiform face area. Further, a higher BIS score was associated with slightly better performance in face recognition. These findings provide novel evidence that trait motivation, particularly behavioral inhibition, is linked to both the structure and function of the face processing system. This underlines the intrinsic coupling between motivational and perceptual systems, blurring the presumed divide between affective and perceptual processes.

## 1 Introduction

It has been increasingly recognized that the mind is highly integrative, with functional modules dynamically collaborating to support an animal’s adaptive interactions with the environment (Bernard et al., 2005; LeDoux, 2012; Russell, 2003; Withagen, 2018). These modules—spanning cognitive, affective-motivational systems, cognitive control, and sensorimotor processes—interact to shape both experience and behavior (Barrett & Bar, 2009; Pessoa, 2008). From this perspective, individual differences in traits may reflect distinct strategies for adapting to environmental demands and opportunities (Gray, 1982; Matthews, 2008; Matthews, 2018), and it is possible that all stages of information processing, from early perceptual encoding to affective evaluation and response execution, may show coordinated idiosyncrasies that together support a coherent and adaptive strategy in interpreting and responding to the environment.

Indeed, long-term individual variations in motivation, often conceptualized as trait motivation, have been shown to impact how individuals process rewarding or threatening stimuli (Elliot, 2013). Research on the influence of trait motivation on the processing of reward- and threat-related stimuli suggested that individuals with different motivational dispositions perceive and respond to their environment in distinct ways. Individuals with stronger approach motivation tend to exhibit heightened sensitivity to rewarding stimuli, such as positive facial expressions, appetitive food cues, and monetary gains, with enhanced neural responses in reward-related regions like the ventral striatum and orbitofrontal cortex (Kennis et al., 2013; Standen et al., 2022). Conversely, individuals with stronger avoidance motivation display increased sensitivity to threats, responding more strongly to negative stimuli such as angry faces or potential punishments, often showing heightened activation in regions such as the amygdala and anterior cingulate cortex (Kennis et al., 2013; Standen et al., 2022). These findings indicate that trait motivation modulates perception when stimuli carry explicit emotional or motivational significance. However, it remains unclear whether these trait-like motivational dispositions extend their influence beyond explicit reward or threat processing to shape more general perceptual mechanisms. Does trait motivation bias the processing of neutral stimuli, such as faces with neutral expressions or general objects? If motivational traits are truly ingrained in perceptual and cognitive processing, their impact may not be limited to emotionally salient stimuli but could also shape the general perceptual encoding preposition and corresponding neural correlates.

To test this possibility, the present study takes face processing as a specimen of general perceptual processing and asks whether trait motivation influences the structure and function of the face-selective regions in the ventral visual pathway, including the fusiform face area (FFA), occipital face area (OFA), and superior temporal sulcus (STS) (Haxby et al., 2000; Kanwisher et al., 1997). Faces, as visual stimuli, hold immense ecological significance in human cognition. The ability to rapidly detect and recognize faces allows individuals to respond adaptively to threats or affiliative cues (Darwin, 1872; Leopold & Rhodes, 2010). Given this evolutionary importance, it is unsurprising if face perception is modulated by emotional and motivational propositions, making it a plausible candidate for modulation by trait motivation. In the present study, trait motivation was operationalized as a BAS/BIS score, measured according to Gray’s Reinforcement Sensitivity Theory (Gray, 1982). RST is a neurophysiological model of personality that explains individual differences in motivation through two primary neurobehavioral systems: the Behavioral Activation System (BAS) and the Behavioral Inhibition System (BIS). The original version of RST (Corr et al., 1995; Gray, 1982) posits that the BAS governs responses to rewarding stimuli, facilitating approach behaviors toward appetitive stimuli. In contrast, the BIS is sensitive to punishment, non-reward, and threats, promoting avoidance behaviors to escape aversive stimuli. Previous studies have examined the effect of BAS and BIS systems on emotional face processing. Individuals with low BAS and high BIS show heightened accuracy in recognizing fearful faces among different emotional faces (Montagne et al., 2005), whereas high BAS individuals exhibit greater vigilance toward angry faces compared to high BIS individuals (Putman et al., 2004). Furthermore, BAS is associated with enhanced left-hemispheric EEG activity in response to positive faces, while BIS is linked to increased right-hemispheric EEG activity when processing negative faces (Balconi & Mazza, 2010). However, it remained unclear whether the individual difference in the BAS/BIS system is associated with individual differences in the neural and behavioral properties of face processing in more general, non-emotional contexts.

In the present study, we measured the ROI-wise gray matter volume in bilateral FFA, OFA, and STS based on structural MRI data, and examined their relation with BIS and BAS scores. To foresee our findings, we observed a significant correlation between the GMV of bilateral FFA and BIS. This correlation was replicated by voxel-wise analysis across the whole brain. Furthermore, we examined whether the association with trait motivation is also reflected in face recognition performance in an old-new task.

## 2 Method

### 2.1 Participant

A total of 264 participants (125 males, 139 females) were recruited for the study, with a mean age of 21.754 years (SD = 1.015, range: 18–25). Eighteen participants were excluded for not completing the functional imaging experiment, and another seven were excluded due to incomplete participation in the behavioral experiment. All analyses were based on the maximum available sample sizes: VBM (n = 264), functional activity (n = 246), and behavioral experiment (n = 239). The study was approved by the Ethics Committee of Beijing Normal University. Written informed consent was obtained from all participants prior to the experiment.

### 2.2 Behavioral Activation System and Behavioral Inhibition System Measurements

A validated Chinese version of the BAS/BIS scale (Li et al., 2008) was used to assess BAS/BIS. The BAS/BIS scale consists of thirteen BAS items and seven BIS items (Carver & White, 1994). Following Li et al. (2008), we excluded items 1 and 18 due to their low correlations with other items (r < 0.3) and ambiguous factor loadings. The final scale comprised thirteen BAS items and five BIS items. Higher BAS scores indicate greater approach trait motivation and heightened sensitivity to reward signals, whereas higher BIS scores indicate greater avoidance trait motivation and heightened sensitivity to punishment signals. BAS and BIS scores were calculated as the sum of their respective item scores. Internal consistency for the scales was acceptable, with Cronbach’s α = 0.761 for BAS and Cronbach’s α = 0.738 for BIS. Due to the moderate correlation between BAS and BIS total scores (*r* = 0.274, *p* < 0.001), we calculated residual BIS and BAS scores by regressing BAS from BIS and vice versa, unless otherwise specified (Ide et al., 2020).

### 2.3 Image acquisition

Scanning was conducted at BNU Imaging Center for Brain Research, Beijing, China, on a Siemens 3T scanner (MAGENTOM Trio, a Tim system) with a 12-channel phased-array head coil. Structural T1-weighted images were acquired with a 3D magnetization-prepared rapid acquisition gradient-echo (MP-RAGE) scan (image = 128 sagittal slices, TR/TE/TI = 2530/3.39/1100 ms, flip angle = 7°, FOV = 256×256 mm^2^. Voxel size: 1×1×1.33 mm^3^). They were collected for voxel-based morphometry analysis, registration, and anatomically localizing the functional activations. Functional blood-oxygen-level-dependent images were acquired with a T2*weighted gradient-echo, echo-planar-imaging (GRE-EPI) sequence (TR/TE = 2000/30 ms; flip angle = 90°, in-plane resolution = 3.1 × 3.1 mm). Whole-brain coverage for the functional data was obtained using 30 contiguous interleaved 4.8 mm axial slices.

### 2.4 Functional localizer for the face-selective regions

A dynamic face localizer was used to define the inferior temporal face-selective regions (Pitcher et al., 2011). Specifically, three block-design functional runs were conducted with each participant. Each run contained two block sets, intermixed with three 18 s fixation blocks at the beginning, middle, and end of the run. Each block set consisted of four blocks with four stimulus categories (faces, objects, scenes, and scrambled objects), with each stimulus category presented in an 18 s block that contained six 3 s movie clips. During the scanning, participants were instructed to passively view the movie clips (for more details on the paradigm, see Pitcher et al., 2011). Only neural activity in face-viewing and object-viewing conditions were analyzed in the present study, to calculate the face selectivity of specific brain regions (i.e., faces versus objects).

### 2.5 VBM data preprocessing

Voxel-based morphometry (VBM) analysis was performed using SPM8 (Statistical Parametric Mapping, Wellcome Department of Imaging Neuroscience, London, UK) and DARTEL (Wellcome Department of Imaging Neuroscience) to quantify gray matter volume at each voxel. Image quality was first assessed by visual examination. The origin of each brain was then manually set to the anterior commissure for each participant. Third, images were segmented into gray matter (GM), white matter, cerebrospinal fluid, and everything else (e.g., skull and scalp) using a unified segmentation approach (Ashburner & Friston, 2005). The segmented GM images were rigidly aligned and resampled to 2 × 2 × 2 mm and then nonlinearly registered with DARTEL, which involves iteratively computing a study-specific template based on the tissue maps from all participants and then warping all participants’ GM images into the generated template to improve the alignment (Ashburner, 2007). The resulting GM images were normalized to standard MNI space, and the GM voxel values were modulated by multiplying the Jacobian determinants derived from the registration to preserve the volume of tissue from each structure (Good et al. 2001). The modulated GM images were then smoothed with an 8-mm full width at half maximum (FWHM) isotropic Gaussian kernel. Finally, to further exclude non-gray matter voxels, the modulated images were masked using the MNI152 gray matter mask with a probability threshold of 0.2.

### 2.6 fMRI data processing

fMRI data were preprocessed using tools from the FMRIB Software Library (FSL, http://www.fmrib.ox.ac.uk/fsl) and in-house Python tools. The preprocessing steps included high-pass temporal filtering (120-second cutoff), motion correction, brain extraction, spatial smoothing (Gaussian kernel with FWHM = 6 mm), and grand-mean intensity normalization. Statistical analysis of the time series was performed using FILM (FMRIB’s Improved Linear Model) with local autocorrelation correction.

First-level analyses were conducted independently for each run and each session. The Generalized linear model (GLM) included explanatory variables (EVs) representing scenes, faces, objects, and scrambled objects, convolved with a hemodynamic response function. For each EV, the onset times and durations of each stimulus were modeled. To improve model sensitivity, the temporal derivative of each EV was also included. Additionally, six motion correction parameters were incorporated into the model as nuisance regressors to account for residual head motion. We computed face – object contrast in the first-level analysis. After first-level analyses, data from all runs within each session were combined for second-level analyses. Specifically, parameter (beta) images from the first-level analyses were coregistered to each participant’s structural image using FLIRT (FMRIB’s Linear Image Registration Tool) with six degrees of freedom, followed by nonlinear registration to the MNI152 template using FNIRT (FMRIB’s Nonlinear Image Registration Tool) with default parameters. The spatially normalized parameter images, resampled to 2-mm isotropic voxels, were combined across runs within each session using a fixed-effects model.

### 2.7 Region of interest analysis

We defined three bilateral regions of interest (ROIs): the fusiform face area (FFA), the occipital face area (OFA), and the posterior superior temporal sulcus (pSTS). Each bilateral ROI was composed of the top 100 voxels with the highest probability from the corresponding probabilistic activation map of FFA, OFA, and pSTS in Brain Activity Atlas (Zhen et al., 2015) in each hemisphere, resulting in 200 voxels per ROI. The peak probabilistic selectivity voxels were located at MNI coordinates (−40, −52, −22) and (42, −50, −22) for the right and the left hemisphere, for the FFA; (−42, −80, −16) and (42, −80, −16) for the OFA; and (−56, −42, 6) and (48, −38, 4) for the pSTS, respectively.

To identify brain regions whose mean gray matter volume (GMV) was associated with trait motivation, we conducted partial correlation analyses between the residual scores of BIS or BAS and the mean GMV of each ROI, controlling for total gray matter volume, age, and sex (*p* < 0.05, Bonferroni corrected).

### 2.8 Voxel-wise analysis

Beyond the ROI-based analysis, we conducted voxel-wise whole-brain searchlight analysis to identify regions where BIS residuals were significantly correlated with gray matter volume across participants. Age (in years), total gray matter volume, and sex were included as covariates. A voxel-level threshold of *p* < 0.001 (uncorrected) was applied in line with previous VBM studies examining correlations between personality traits and gray matter morphometry (Schilling et al., 2013; Van Schuerbeek et al., 2011), along with a minimum cluster size threshold of 50 voxels.

We further assessed the face selectivity of the identified clusters by examining whether each cluster demonstrated greater activation for faces compared to objects in the functional localizer scan. Specifically, we tested the contrast of faces > objects in these clusters at the group level, using a one-tailed threshold of *p* < 0.05 (Bonferroni corrected).

### 2.9 Behavioral task

The old/new recognition memory paradigm was used to measure participants’ face recognition performance. The face recognition task used 40 face images and 40 flower images. The face images were from an in-house database, depicting adult Chinese faces in grayscale, with the external contours removed (leaving an oval shape, not showing the hair and the neck, Figure 1c). The flower images were grayscale pictures of flowers with leaves and backgrounds removed. Flowers were included as a control to measure general memory ability, allowing the removal of general memory influences on face-specific memory performance through regression analysis. The task consisted of two blocks: a face block and a flower block. Each block consisted of a learning phase and a testing phase. During the learning phase, 20 images of that category were presented for 1 second each, with an inter-stimulus interval (ISI) of 0.5 seconds. These 20 images were presented twice in a loop. In the testing phase, 10 learned images were randomly intermixed with 20 new images of the same category, and each image was presented twice. While viewing each image, participants were required to determine whether the image had been shown during the learning phase (Li et al., 2010). The face-specific recognition accuracy was calculated as the residual after regressing general memory accuracy (measured by flower recognition) from face recognition accuracy.

**Figure 1.**
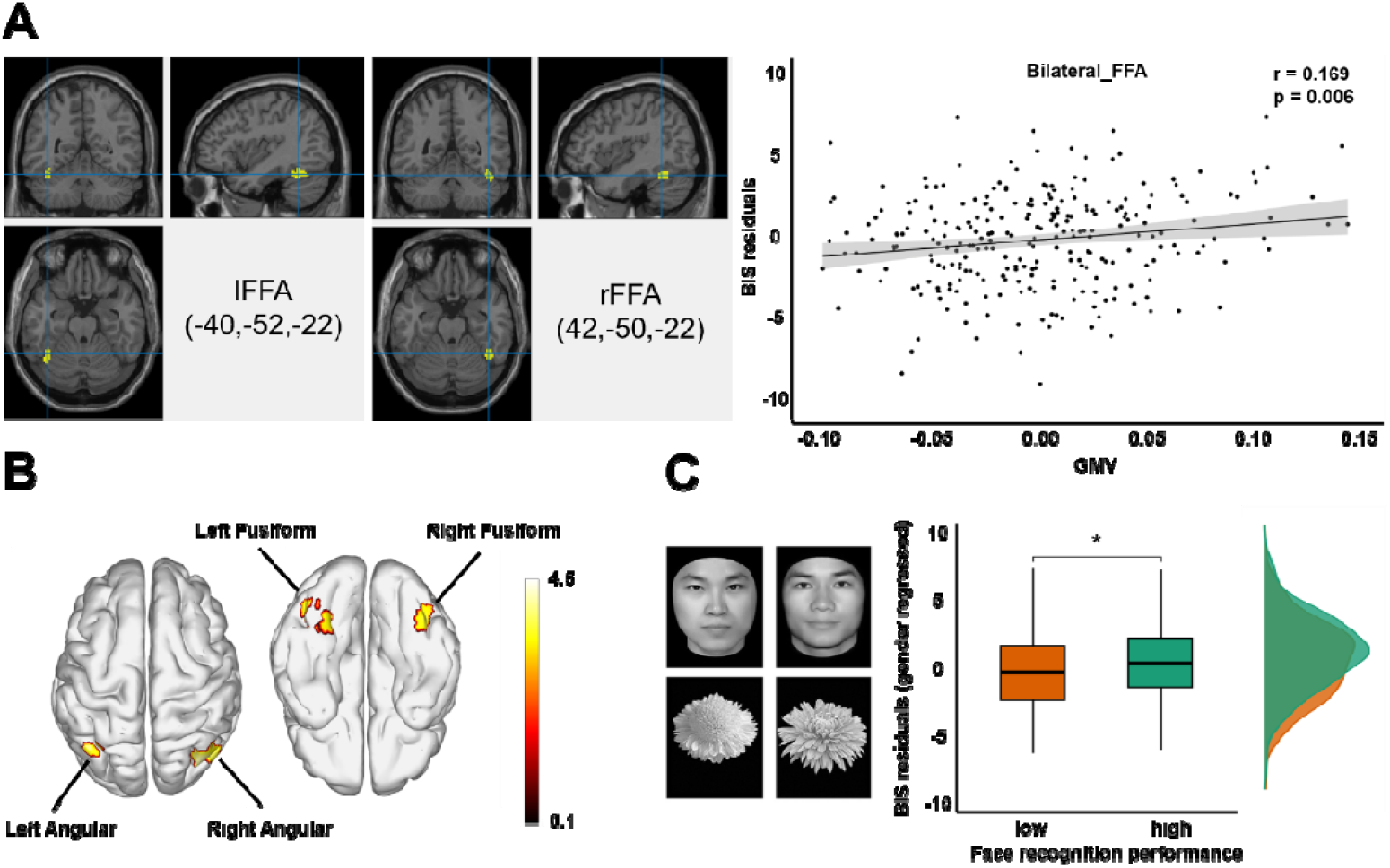
BIS-residual correlates with GMV in FFA and face recognition performance. (A) Left: Anatomical localization of the FFA ROI in the left and the right hemisphere, with the MNI coordinates of their corresponding centers indicated. Right: the scatter plots depicting the partial correlations between BIS residual scores and GMV in these ROIs (adjusted for BAS scores, total gray matter volume, age, and sex). (B) Voxel-wise analysis results showing significant clusters where BIS scores were positively correlated with GMV (voxel-level *p* < 0.001, uncorrected; cluster size > 50 voxels). The color scale represents T-values from the regression analysis. (C) Left: Example face and flower stimuli of the old–new recognition task. Right: a boxplot with probability density overlays comparing BIS residuals (adjusted for general memory performance, BAS scores, age and sex) between participants with high and low face recognition performance. Participants with higher face recognition performance also tend to have higher BIS levels. rFFA: Right fusiform face area. lFFA: Left fusiform face area. GMV: Gray matter volume.

### 2.10 Behavioral analysis

To investigate whether BIS has behavioral implications on face processing, we compared participants’ BIS index with high and low face-specific recognition performance. Participants were divided into two groups based on their face-specific memory performance, with the top 50% categorized as high face memory performers and the bottom 50% as low face memory performers. An independent-sample *t*-test was performed to compare BIS scores between the two groups after controlling BAS scores, sex, age, and general memory performance.

## 3 Results

### 3.1 BIS Residual correlates with ROI-Wise GMV in FFA

We first investigated the relationship between trait motivation and the gray matter volume of regions in the core face network. We examined three bilateral ROIs—the fusiform face area (FFA), occipital face area (OFA), and posterior superior temporal sulcus (pSTS). They were defined as the top 100 highest-probability voxels in each hemisphere, based on the corresponding probabilistic atlas (Zhen et al., 2015). For each ROI, we conducted partial correlation analyses between the ROI’s mean GMV and the BIS or BAS residuals, respectively, with total gray matter volume, age, and sex included as covariates. We observed a significant positive partial correlation between BIS residual scores and mean gray matter volume (GMV) in the bilateral FFA (bilateral FFA: *r*_*partial*_ = 0.169, *p* = 0.006, Figure 1a), indicating that higher BIS levels are associated with larger GMV in bilateral FFA. The partial correlations between mean GMV of other ROIs and BIS residual were not significant, nor were the GMV-BAS partial correlations (Table 1). This pattern remained consistent when the bilateral FFA ROI was divided into left and right hemispheric regions (Supplementary Materials S1). The association was also held under alternative FFA definitions (Supplementary Materials S2).

**Table 1.** Partial correlation coefficients between ROI-wise GMV and BIS/BAS residuals.

| ROI | BIS | BAS |
| --- | --- | --- |
| Bilateral FFA | .169 (.006) | -.007 (.913) |
| Bilateral OFA | .066 (.285) | .026 (.676) |
| Bilateral pSTS | .088 (.156) | -.019 (.758) |

*Note.* Partial correlations were conditioned on sex, age (in years), and total GMV. Values in parentheses indicate the corresponding $p$ -values.

### 3.2 BIS residual correlates with voxel-wise GMV in face-selective clusters in fusiform area

In addition to the ROI analysis, we searched for any voxels in the whole brain that showed a significant correlation between BIS residuals and voxel-wise gray matter volume across participants, with age (in years), total gray matter volume, and sex as covariates. The analysis revealed significant positive GMV-BIS associations in four clusters in the bilateral fusiform gyrus and bilateral angular gyrus (*p* < 0.001, uncorrected, Table 2), and no significant negative GMV-BIS associations were observed (Figure 1C). Among them, the only cluster that showed significant face-selective activation was in the right fusiform gyrus (*t* (245) = 8.946, *p* < 0.001). This cluster overlapped with our FFA ROI with 13 voxels in the right hemisphere. In contrast, the cluster in the left fusiform gyrus exhibited opposite categorical selectivity (*t* (245) = −9.212, *p* < 0.001) and showed little overlap with our FFA ROI in the left hemisphere (n = 2). The cluster in the right angular gyrus exhibited selectivity for the non-face category (*t* (245) = −9.684, *p* < 0.001), whereas the left angular gyrus showed no significant selectivity (*p* = 0.711).

**Table 2.** Clusters that showed significant associations between GMV and BIS residuals (voxel-wise p < 0.001, uncorrected, cluster size > 50 voxels).

| Cluster | Cluster size | Peak MNI coordinate (x, y, z) |
| --- | --- | --- |
| Right Fusiform | 58 | (40, -54, -14) |
| Left Fusiform | 146 | (-36, -56, -10) |
| Right Angular | 308 | (42, -74, 38) |
| Left Angular | 92 | (-38, -68, 52) |

Convergently, the results of the ROI analysis and voxel-wise analysis suggest a significant positive association between BIS and gray matter volume in the FFA, particularly in the right hemisphere.

### 3.3 High performance in face recognition is associated with high BIS

To further investigate whether BIS is associated with face recognition performance, we measured face recognition with an old-new recognition task and examined its association with BIS. Face recognition performance was divided into high and low groups based on the median split. We observed a significant difference between the BIS scores of individuals with high and low face recognition performance after controlling for BAS scores, sex, age, and general memory performance (*t* (237) = −2.060, *p* = 0.040, Cohen’s *d* = −0.257; Figure 1C), suggesting that individuals with higher face recognition performance also tend to have higher BIS levels.

## 4 Discussion

The present study reported a positive association between BIS score and gray matter volume (GMV) in the FFA, a key brain region involved in face processing. Supporting the neural findings, we further observed that individuals with high BIS tended to exhibit better face recognition performance. Together, these findings suggested that individual difference in trait motivation is associated with individual differences in face processing, illustrating a potential link between trait motivation and face perception.

In our study, ROI-wise and whole-brain voxel-wise analysis convergently suggested that participants with higher BIS, that is, those with a stronger tendency for increased attention and passive avoidance in response to perceived threats or punishment, had greater GMV in the FFA, particularly in the right hemisphere. It is well established that the FFA, particularly the right FFA, is crucial in high-level face perception, including processing invariant features of faces and extracting the global configuration of facial features (Halgren et al., 1999; Haxby et al., 1999; Kanwisher et al., 1997 & Meng et al., 2012), and plays a particularly essential role in face recognition (Duchaine & Yovel, 2015). In contrast, OFA was considered a lower-level region in the processing stream of faces, primarily specialized in processing the structural components of faces, such as individual facial features (eyes, nose, mouth, Haxby et al., 2000). The STS, on the other hand, has been associated with the processing of dynamic facial information, including gaze direction, emotional expressions, and facial muscle movement in communication (Haxby et al., 2000). The FFA-specific effect observed in our study suggested that trait motivation, particularly the BIS system, may be selectively associated with the higher-level processes in face perception, rather than the early-stage structural encoding of facial features or the dynamic and contextual aspects of face processing, such as the processing of transient emotional expressions and social signals.

Notably, the link between BIS and face recognition is also manifested at the behavioral level. Our results indicated that individuals with better face recognition performance tend to have high BIS levels. We speculate that this is because the long-term motivational valence of a specific face largely depends on its identity. For instance, the valence of a face can be decided by its identity—whether it belongs to a stranger, who may pose a potential threat, or a recognized kin, who is typically non-threatening. Trait motivation might have been ingrained in this identity-dependent social evaluation process, with individuals who are more sensitive to threats (i.e. those with higher BIS) allocating greater neural resources in face recognition than those who are less sensitive. Another possibility is that this BIS-face perception association is based on affective processing. Previous studies have suggested that individuals with anxiety often misinterpret neutral expressions as threatening (Phillips et al., 1997) and that high BIS individuals are more likely to perceive hostility in any facial expression (Knyazev et al., 2008). In the current study, it is possible that high BIS participants perceived neutral faces as threatening stimuli, leading to increased neural investment in face processing, and therefore greater GMV in the right FFA, compared to low BIS participants. Future study is needed to directly test these speculations.

Admittedly, previous studies have not reported any association between the structural features of FFA or other regions in the face network and trait motivation. This is possible because they mostly conducted whole-brain analyses (Fuentes et al., 2012; Ide et al., 2020), which may have failed to capture the small effect sizes captured by our hypothesis-driven analysis and larger sample size. Additionally, different from previous studies of Western cultural context and either earlier (Ide et al., 2020) or later (Cherbuin et al., 2008) developmental stages, our participants were uniformly college students in their early adulthood in a Chinese cultural context. In Chinese culture, maintaining long-term relationships and respecting social roles are deeply ingrained values, especially for individuals navigating crucial life transitions typically in early adulthood. This cultural context might bestow extra motivational relevance to face recognition, and lead to a particularly strong association between trait motivation and face recognition in our sample. Future research is needed to test the cross-cultural generality of our findings and to further identify the cultural and developmental modulators of the motivation-perception connection.

Another slightly surprising aspect of our results was that our voxel-wise analysis did not replicate findings of the association between BIS and brain regions proposed to be neurobiological correlate of BIS (Firth et al., 2024, Gray, 1982), nor brain regions typically implicated in affective-motivational processes. Also, we did not find evidence for any association between face-selective regions and BAS. These could be due to the small effect size of the structural effect of trait motivation in these regions, or the cultural or developmental-specific factors which, as we discussed earlier, might have biased the association between perception and trait motivation from a dynamic and affective route to a relatively passive, perceptual route. Future study is required to directly test these possibilities.

Though we took face perception as a specimen, we do not claim that the observed association with trait motivation is specific to face perception. Instead, some clusters identified in the whole-brain searchlight analysis did not exhibit face selectivity, suggesting that the association between trait motivation and perception may extend beyond face perception. Consistent with this speculation, the brain regions we found to be associated with BIS outside the face-selective areas primarily emerged in higher-level visual regions, in the left fusiform gyrus and bilateral angular gyrus. The fusiform gyrus is established for processing visual information related to object recognition and categorization (Joseph, 2001). The angular gyrus, on the other hand, is involved in integrating sensory information from various modalities and plays a significant role in semantic processing, memory retrieval, and spatial awareness. Both regions are integral to the processing of complex visual stimuli and contribute to our ability to interpret and respond to the environment. The association between these regions and BIS suggests that the influence of trait motivation, particularly the behavioral inhibition system (BIS), may be generalizable to broader perceptual processes. Future research should investigate the category specificity of this BIS-perception association, which may help reveal the origin of this link.

Overall, consistent with recent perspectives that view affective-motivational processes as integral in adaptively interpreting and responding to the environment (Bernard et al., 2005; LeDoux, 2012; Russell, 2003; Withagen, 2018), our findings provided novel empirical evidence suggesting an association between trait motivation and perception. Neurobiological models of personality, including RST, have long emphasized the role of neurobiological factors, including genes and neurotransmitter systems, in shaping individual differences and long-term behavioral tendencies. These factors, rather than being confined to a specific functional module, operate across the central nervous system, likely to generate broad, domain-general impacts and contribute to co-variation across functional modules. In line with this idea, prior research has indicated that sensitivity to stimuli might be one of the fundamental elements underlying personality differences (Wolf et al., 2008). In the case of avoidance motivation, enhanced face perception, and possibly object perception in general, may be adaptive when paired with high BIS levels, because it may facilitate threat detection and conflict monitoring. However, it is important to note that the current study provides solely correlational evidence, without exploring the underlying mechanism, and the causal relationship between BIS and face processing remains unconfirmed. Future research is needed here, probably in the form of longitudinal studies or leveraging genetic neurochemical approaches.

## CRediT authorship contribution statement

Hangshek Lau: Writing – original draft, Writing – review & editing, Visualization, Formal analysis. Yiying Song: Writing – review & editing. Shan Xu: Writing – review & editing, Conceptualization, Funding acquisition. Jia Liu: Writing – review & editing, Conceptualization.

## Acknowledgments

Shan Xu discloses support for the research of this work from the National Natural Science Foundation of China (32371099), and Jia Liu discloses support for the research of this work from the National Natural Science Foundation of China (T248810018), the Beijing Municipal Science & Technology Commission, and the Administrative Commission of Zhongguancun Science Park (Z221100002722012).

## Data availability statement

The data underlying this study can be obtained from the corresponding author, Shan Xu, upon reasonable request. To protect participant privacy, the data are not publicly available.

