## Supplementary Materials S1& Supplementary Materials S2 for "Trait motivation is associated with Fusiform face area Morphometry: Evidence from a Chinese Youth Sample"

**The association between FFA GMV and BIS residual scores remained consistent when the bilateral FFA ROI was divided into left and right hemispheric regions.**

We examined the left and right FFA separately. The pattern of partial correlations between BIS and FFA GMV remained significant, consistent with the main findings (see Supplementary Table 1).

Supplementary Table 1. Partial correlation coefficients between GMV in the two FFAs defined by probabilistic activation maps and BIS residuals.

| Probabilistic activation map-based ROIs | BIS |
| --- | --- |
| rFFA | .168 (.007) |
| lFFA | .137 (.027) |
| *Note.*  Partial correlations were conditioned on sex, age (in years), and total GMV. Values in parentheses indicate the corresponding p-values. | |

**Supplementary Materials S2**

**The association between FFA GMV and BIS residual scores remained consistent across different ROI definitions.**

We examined the robustness of the association between BIS residual scores and FFA gray matter volume (GMV) using three bilateral FFA definitions other than the one used in the main text (Zhen et al., 2015). FFA was defined based on the average bilateral FFA coordinates reported in a meta-review of face-related fMRI studies (Berman et al., 2010) and the peak face-selective activation coordinates in FFA observed in two representative paradigms (Fox et al., 2009; Kanwisher et al. 1997). In each case, GMV was extracted using two 6-mm radius spherical masks centered at the reported MNI coordinates for the left and the right FFA. The **Berman ROI** was centered at (−42, −57, −17) and (43, −55, −18); the **Kanwisher ROI** at (−36, −63, −17) and (41, −54, −18); and the **Fox ROI** at (−40, −42, −28) and (38, −47, −29). As in the main analysis, partial correlations between BIS residual scores and GMV in each ROI were computed, controlling for sex, age (in years), and total GMV. A significant positive partial correlation between BIS residual scores and mean GMV was observed in all cases. The results were presented in Supplementary Table 2.

Supplementary Table 2 Partial correlation coefficients between GMV in three FFAs (defined using 6-mm radius spheres) and BIS residual scores.

| ROI | Pearson’s *r* |
| --- | --- |
| The **Berman ROI** | .209 (<.001) |
| The **Kanwisher ROI** | .233 (<.001) |
| The **Fox ROI** | .127 (.040) |
| *Note.*  Partial correlations were conditioned on sex, age (in years), and total GMV. Values in parentheses indicate the corresponding p-values. | |
